# Phylogenomic relationships and gene content of four forms of the *Culex pipiens* (Diptera: Culicidae) complex

**DOI:** 10.64898/2026.07.25.740720

**Authors:** Tyler Maire, Erin Maley, Theresa Miller, Kyle Kosinski

## Abstract

The *Culex pipiens* L. (Diptera: Culicidae) complex includes the main vectors of West Nile virus in temperate regions, yet few studies have compared the genomes of its members. We compared chromosome-scale assemblies of *Cx. pipiens* form *pipiens*, *Cx. pipiens* form *molestus* Forskål, *Cx. pipiens pallens* Coquillett and *Culex quinquefasciatus* Say, using gene models transferred from the *Cx. quinquefasciatus* annotation and rooting the tree with *Culex perexiguus* Theobald, a species outside the complex. The rooted tree paired *molestus* with *pipiens* and *pallens* with *quinquefasciatus*. Only 59% of the four taxon gene trees agreed; of those that disagreed, 1,746 paired *molestus* with *quinquefasciatus* and *pallens* with *pipiens*, against 1,384 for the other arrangement. Among the best-supported trees, the count was 349 to 184, and loci with intact gene models leaned the same way. In ABBA-BABA tests, *molestus* shared more derived alleles with *quinquefasciatus* than *pipiens* did, and *pallens* shared more with both forms of *Cx. pipiens* than *quinquefasciatus* did, consistent with gene flow between the lineages and *Cx. pipiens* ancestry in *pallens*. Transferred models that shared an error produced false shared derived alleles, which moved the root and inflated two tests until we set such loci aside. Most apparent gene absences were the same gene clustered elsewhere, and gene family changes tracked copy loss during transfer. Median protein identity between forms was near 99%, and synteny was conserved. Comparing genomes in this complex will require independent annotation of each genome, plus checks for errors the annotations share.

## Introduction

Mosquitoes of the *Culex pipiens* L. (Diptera: Culicidae) complex are the main vectors of West Nile virus across temperate North America and Europe, and *Culex quinquefasciatus* Say also transmits St. Louis encephalitis virus and the nematodes that cause lymphatic filariasis (Hayes et al. 2005, Arensburger et al. 2010, Farajollahi et al. 2011). The complex is taxonomically difficult. Its members look alike and hybridize where they meet, and authors have treated them as species, subspecies, or ecological forms (Harbach 2012, Aardema et al. 2022). In current usage, four species are recognized: *Cx. pipiens* and *Cx. quinquefasciatus*, both now found worldwide, and the Australian *Culex australicus* Dobrotworsky and Drummond and *Culex globocoxitus* Dobrotworsky. It also includes the East Asian subspecies *Cx. pipiens pallens* Coquillett and two ecologically distinct forms of *Cx. pipiens*, form *pipiens* and form *molestus* Forskål (Aardema et al. 2022). We compare four of these taxa and call them simply *pipiens*, *molestus*, *pallens,* and *quinquefasciatus* from here on.

The forms differ in ways that matter for transmission. Form *pipiens* is anautogenous, feeds mainly on birds, and needs open space to mate. Form *molestus* is autogenous. It mates in confined spaces and readily bites mammals (Vinogradova 2000, Fonseca et al. 2004). Hybridization between the bird-biting and mammal-biting forms has been proposed to make *Cx. pipiens* an efficient bridge vector, carrying West Nile virus from birds to humans (Fonseca et al. 2004, Farajollahi et al. 2011). The origin of *molestus* has been debated for decades (Haba and McBride 2022). Population genomic data now indicate that it adapted to human environments more than 1,000 years ago, in the Mediterranean or the Middle East (Haba et al. 2025). *Culex pipiens* and *Cx. quinquefasciatus* also hybridize commonly where their ranges meet, though the extent varies by region (Cornel et al. 2003, Kothera et al. 2009).

The status of *pallens* is the least settled. Every *pallens* male sampled in Japan, Korea, and China carried a *Cx. quinquefasciatus* allele at the acetylcholinesterase 2 locus, suggesting sex-linked, asymmetric introgression from *Cx. quinquefasciatus* (Fonseca et al. 2009). A later global genomic analysis went further and concluded that *pallens* most likely arose from past hybridization between *Cx. quinquefasciatus* and *Cx. pipiens* (form *pipiens* or form *molestus*), not from ongoing hybridization (Aardema et al. 2020). Much of this work rested on a handful of markers and on diagnostic assays built to tell the forms apart (Smith and Fonseca 2004, Bahnck and Fonseca 2006, Fonseca et al. 2009). No one has yet examined how the four forms are related using thousands of loci from reference-quality genomes.

That is now possible. Chromosome-scale assemblies exist for all four forms (Liu et al. 2023, Ryazansky et al. 2024, Hesson et al. 2025) and for *Culex perexiguus* Theobald (Ruiz-López et al. 2026), a member of the Univittatus Subgroup of the Pipiens Group that sits outside the complex (Harbach 2011). These genomes open up two kinds of comparison.

The first is phylogenomic. Four taxa allow three unrooted topologies, and if incomplete lineage sorting were the only source of discordance, the two topologies that disagree with the species tree would appear equally often; an excess of one signals introgression (Hibbins and Hahn 2022). An outgroup roots the tree. It also enables ABBA-BABA tests, which ask whether a taxon shares more derived alleles with one of two sister taxa than with the other (Green et al. 2010, Durand et al. 2011). Analyses of this kind uncovered extensive introgression among members of the *Anopheles gambiae* Giles complex (Fontaine et al. 2015). The second comparison is of gene content, the pangenome approach, which sorts groups of orthologous genes into those shared by all members (core), by some (shell), or by only one (cloud) (Tettelin et al. 2005, Golicz et al. 2020, Bayer et al. 2020).

Gene content comparisons are only as good as the annotations behind them. The *Cx. quinquefasciatus* reference annotation is the most complete in the complex. Native annotations of the other forms, where they exist, were made at different times from different evidence and methods. Mixing annotations produced by different methods inflates the apparent number of lineage-specific genes (Weisman et al. 2022), and errors in draft assemblies and their annotations distort gene numbers and gene family sizes (Denton et al. 2014). Transferring one reference annotation to every genome, for example with Liftoff (Shumate and Salzberg 2021), gives a single consistent source. It also brings its own errors.

Here we compare the four forms, rooted with *Cx. perexiguus*, using gene models transferred from *Cx. quinquefasciatus* reference. We ask four questions. How are the forms related, and where does the root fall among them? Is gene tree discordance symmetric, as incomplete lineage sorting predicts, or asymmetric, as gene flow would make it, and which lineages share excess derived alleles? How divergent are the forms? Which apparent differences in gene content, and which phylogenomic signals, survive tests for annotation-transfer artifacts?

## Materials and Methods

### Genome Assemblies

We used chromosome-scale assemblies of *quinquefasciatus* (GCF_015732765.1, VPISU_Cqui_1.0_pri_paternal; Ryazansky et al. 2024), *pallens* (GCF_016801865.2, TS_CPP_V2; Liu et al. 2023), *molestus* (GCA_024516115.1, TS_CPM_V1; Liu et al. 2023), and *pipiens* (GCA_963924435.1, idCulPipi1.1; Hesson et al. 2025). The outgroup was *Cx. perexiguus* assembly idCulPerx1.1 (GCA_964243045.1; Ruiz-López et al. 2026) (Table 1). In each of the five assemblies, three chromosome-scale sequences carry 94.8 to 99.9% of the length. *Culex perexiguus* replaces the outgroup of the preprint of this study, the contig-level assembly of *Culex tarsalis* Coquillett (Main et al. 2021), because most gene models transferred to that assembly were broken.

**Table 1.**
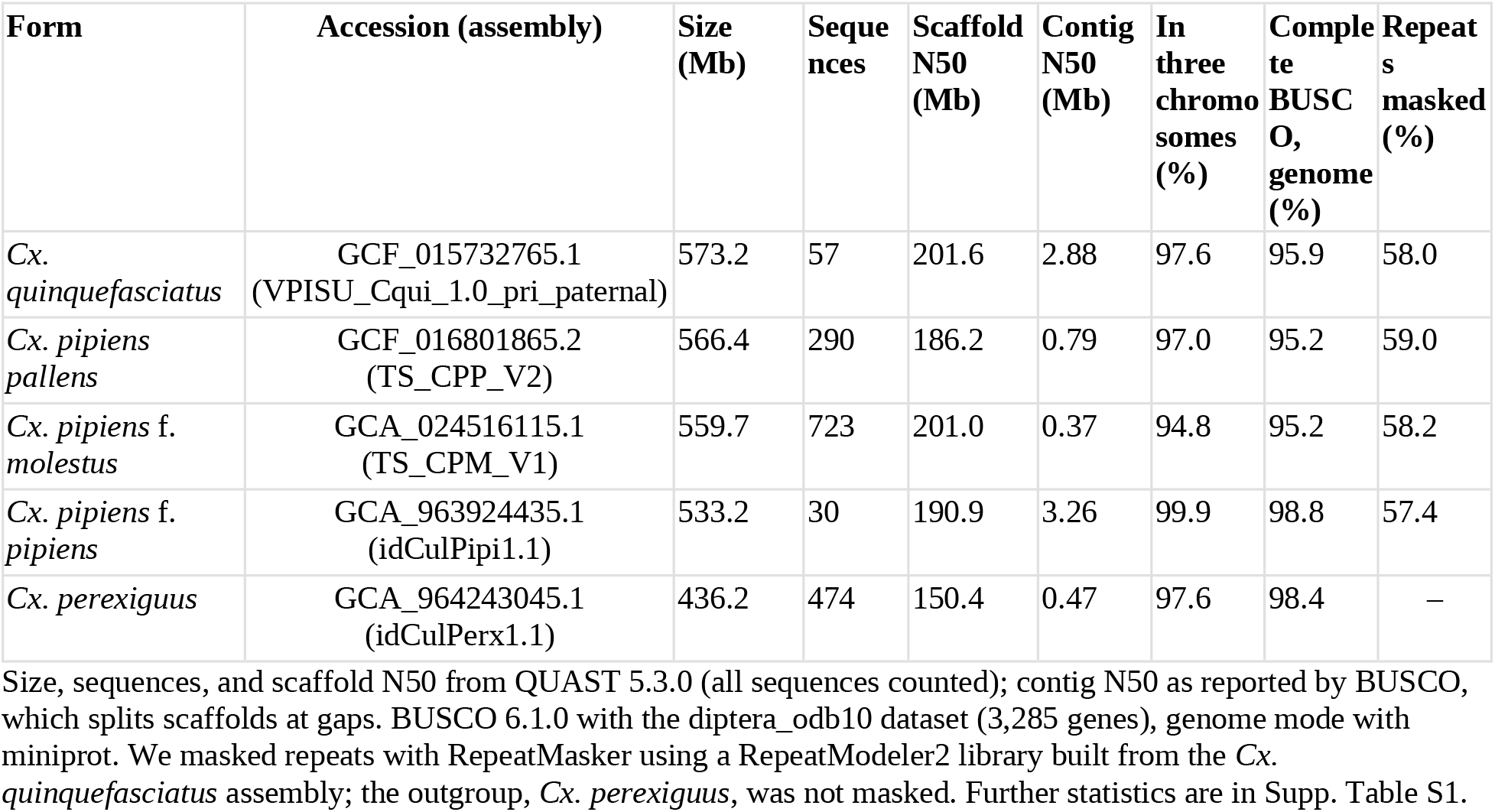
Genome assemblies.

Assembly statistics came from QUAST 5.3.0 (Gurevich et al. 2013), except contig N50, which we took from BUSCO. We assessed completeness with BUSCO 6.1.0 (Tegenfeldt et al. 2025) against the diptera_odb10 dataset (3,285 genes), in genome mode (miniprot 0.18; Li 2023) and in protein mode on the gene sets described below.

### Annotation Transfer and Gene Sets

Gene models from the *Cx. quinquefasciatus* RefSeq annotation (NCBI Annotation Release 100; 17,836 genes, 15,094 protein-coding) were transferred to the other three ingroup assemblies with Liftoff 1.6.3 (Shumate and Salzberg 2021). Liftoff aligns reference genes to the target genome with minimap2 (version 2.24; Li 2018). We required child features such as exons to align with at least 90% identity (-s 0.9) and excluded options that search for extra gene copies (-copies) and repair broken coding sequences (-polish). The *pallens* assembly has a native RefSeq annotation (GCF_016801865.2-RS_2022_12, 16,297 protein-coding genes). Still, we used transferred models there too: mixing a native annotation with transferred ones would turn differences between annotation methods into apparent differences between forms (Weisman et al. 2022). The outgroup was a different case. We kept the Ensembl gene set of *Cx. perexiguus* (12,534 genes; Ruiz-López et al. 2026), because the outgroup served only for orthology and rooting, and because most models we had transferred to *Cx. tarsalis*, the earlier outgroup, were broken.

Liftoff checks every transcript it transfers. It records whether the transcript has a valid open reading frame (ORF), meaning a start codon, a stop codon, and no internal stop codon, and whether its protein matches the reference protein; we summarized these flags for the models kept for analysis. Some reference models are imperfect to begin with. Of the kept reference models, 872 (5.8%) are partial or carry a RefSeq exception for a sequence discrepancy between transcript and genome, and most of their transfers fail the ORF check, whatever the target genome (87 to 89% in the ingroup), so we report ORF failure rates with and without them. We call the four models of a locus intact when the reference model is neither partial nor carries a RefSeq exception, and each transferred model has a valid ORF.

### Orthology and Pangenome Partition

We inferred orthogroups from the five proteomes with OrthoFinder 3.1.5 (Emms et al. 2026), using DIAMOND for similarity searches (Buchfink et al. 2021) and gene trees from multiple sequence alignments (-M msa). We partitioned the orthogroups over the four ingroup forms. Core orthogroups contain genes from all four, shell orthogroups from two or three, and cloud orthogroups from one, while orthogroups holding only *Cx. perexiguus* genes were classed as outgroup only. We did not partition genes that OrthoFinder left out of every orthogroup. Liftoff keeps the reference gene identifier on each transferred model, which let us ask two things of every cloud orthogroup: whether another form had a model of the same gene in a different orthogroup, and whether the orthogroup also contained a *Cx. perexiguus* gene.

### Validation of Gene Absences

Every cloud or shell orthogroup implies that some forms lack the gene. We tested these absences directly. For each absence event, an orthogroup and a form without it, we took the longest ingroup protein of the orthogroup and the genomic locus of the gene encoding it, introns included, and aligned both to the genome of the form lacking it with miniprot 0.18 (--outn 1; Li 2023) and minimap2 2.31 (-x asm20 -N 1; Li 2018). A hit required at least 80% identity and coverage of at least 70% of the protein or 50% of the locus.

We classified events by the first rule that applied. We clustered an event elsewhere when the target proteome contained a model with the query’s gene identifier. We classified it as absent when neither probe found a hit, or weak when hits fell below the thresholds. If a hit passing the thresholds overlapped no protein-coding model, the event was an unannotated locus. If the hit overlapped a model with at least 90% protein identity to the query, the event was clustered elsewhere; identity here meant identical residues over the length of the shorter protein after global alignment, set to 0 when the shorter protein was less than 30% of the longer protein’s length. Because only less similar models overlapped, the event was paralog-only. Only absent and paralog-only events support a real difference in gene content. Clustered elsewhere and unannotated locus events are annotation or clustering artifacts.

### Four Taxon Phylogenomics

We defined single-copy orthologs as orthogroups with exactly one gene in each ingroup. We aligned their proteins with MAFFT (--auto; Katoh and Standley 2013), trimmed them with trimAl (-automated1; Capella-Gutiérrez et al. 2009), and discarded alignments shorter than 50 columns after trimming. We inferred the species tree from the concatenated alignments with IQ-TREE 3.1.3 (Wong et al. 2026) and assessed support with 1,000 ultrafast bootstrap replicates (UFBoot; Hoang et al. 2018). Each locus formed its own partition, and ModelFinder (Kalyaanamoorthy et al. 2017) chose a model for each. For this matrix, ModelFinder considered only equal rates and the +I, +G4, and +I+G4 models of rate heterogeneity (-mrate E, I, G, I+G), because IQ-TREE 3.1.3 stopped with an internal error while fitting FreeRate models. In an earlier run of the workflow on nearly the same loci, with FreeRate models allowed, they were selected for 8 of 9,098 loci (docs/v5_run.md in the repository). Gene trees for each locus used the full model set. We computed gene concordance factors (gCF) from the gene trees and site concordance factors (sCF) from parsimony-informative sites (Minh et al. 2020). Because every sampled quartet includes all four taxa, the sCF is simply the share of parsimony-informative sites supporting each split. This tree is unrooted. We used the outgroup in the analyses described in the next section.

To test whether discordance was symmetric, we counted the gene trees supporting each of the two discordant topologies. We compared them with an exact two-sided binomial test against equal frequencies, with Clopper-Pearson 95% confidence intervals. We then repeated the test on subsets. One series kept gene trees whose internal branch had UFBoot support of at least 50, 70, 80, 90, and 95. Another kept gene trees whose internal branch was longer than IQ-TREE’s minimum length (10^−6^ substitutions per site), since a branch at the minimum carries no inferred substitution. We also split gene trees by whether they had a terminal branch longer than 0.1, because those trees are most exposed to long-branch attraction, and we ran the test on loci with four intact gene models.

We also counted parsimony-informative sites supporting each topology, summed over all loci and over loci with intact models, and, to give each locus one vote, we recorded the topology supported by the most informative sites within each locus (ties and loci without such sites were set aside). An excess of one discordant topology is expected from introgression. It can also arise from ancestral population structure (Hibbins and Hahn 2022, Lanfear and Hahn 2024), and an unrooted quartet test cannot show which pair of lineages exchanged genes.

### Rooted Species Tree and ABBA-BABA Tests

Orthogroups with exactly one gene in each ingroup form and in *Cx. perexiguus* gave 6,846 five-taxon loci. We aligned and trimmed their proteins as above. We then placed the coding sequences on the trimmed protein alignments codon by codon, so that each nucleotide alignment holds the three codon positions of every amino acid column kept after trimming; 6,831 loci had trimmed alignments of at least 50 columns.

Transferred models can share an error. One example is a reading-frame shift where RefSeq corrected the reference model for an error in the reference genome. Every form that carries the error then carries the same misaligned codons, and these look like derived alleles shared by those forms. For that reason, we inferred the rooted tree from the 4,592 loci with intact four-ingroup models, using concatenated protein alignments with one partition per locus, models chosen by ModelFinder, 1,000 UFBoot replicates, and *Cx. perexiguus* as the outgroup. Concordance factors came from the same loci: gCF from their gene trees and sCF from 100 sampled quartets per branch, and we counted rooted gene-tree topologies from the same gene trees. To show the effect of these errors, we also inferred the tree from all loci and counted derived states shared by each combination of forms, polarized by the outgroup, for each locus class (Supp. Table S13).

ABBA-BABA tests (Green et al. 2010, Durand et al. 2011) used the codon alignments. We kept sites where all five taxa carried A, C, G, or T, and the four taxa in a test carried two alleles, taking the outgroup allele as ancestral. For taxa P1 and P2, sisters relative to P3, D = (ABBA − BABA)/(ABBA + BABA), where ABBA sites have the derived allele in P2 and P3 and BABA sites have the derived allele in P1 and P3. Incomplete lineage sorting alone gives D = 0. D < 0 means that P1 and P3 share more derived alleles. Standard errors came from a weighted block jackknife (Busing et al. 1999) over 5 Mb windows of the reference assembly, with each locus placed by the position of its reference gene, and two-sided *P* values from Z = D/SE were Bonferroni-corrected for the four tests.

We fixed these four tests before the analysis, based on the four-taxon species tree: D(*pallens*, *quinquefasciatus*; *pipiens*), D(*pallens*, *quinquefasciatus*; *molestus*), D(*molestus*, *pipiens*; *pallens*), and D(*molestus*, *pipiens*; *quinquefasciatus*). We computed each on all sites and on third codon positions, on all loci, and on loci with intact models. Unless we say otherwise, we report all sites on loci with intact models. As a further check, we filtered loci by the lowest pairwise nucleotide identity among their four ingroup sequences over the codon alignment, at thresholds of 95 and 98%.

### Divergence and Genome Identity

We compared terminal branch lengths between the concatenated four-taxon tree and the per-locus gene trees. We computed pairwise protein identity between forms for each trimmed alignment, over columns where both forms had a residue. For whole genomes, skani (skani dist - s 80; Shaw and Yu 2023) estimated average nucleotide identity (ANI) and aligned fraction on the complete assemblies. We also aligned the three chromosome-scale sequences of each pair of ingroup assemblies with minimap2 2.31 (-x asm20 --secondary=no). We used alignment identity as the length-weighted mean of one minus the divergence that minimap2 recorded for each alignment.

### Gene Family Evolution and the Transfer Control

We modeled changes in gene family size with CAFE 5 (Mendes et al. 2020), using OrthoFinder gene counts for the four ingroup forms in 13,605 orthogroups with ingroup genes (no form exceeded 100 genes) and a gamma model with three rate categories. CAFE tests only families present at the root, so it set aside the 741 orthogroups without genes on one side, leaving 12,864. We used the species tree as the rooted tree, with the outgroup. Divergence times are unknown, so we set terminal branch lengths to 1 and the two branches from the root to 0.5.

CAFE 5 defines the *P* value of a family as the probability, under the fitted model and given the root size, of counts no more likely than those observed, taking the largest value over root sizes from 1 to 1.25 times the family’s largest count. It estimates this from 1,000 simulated families per root size without a fixed random seed, so the estimate varies between runs. Families with the same counts share one estimate, so a count pattern held by hundreds of families crosses a threshold all at once, and in 20 repeated runs on our data with the fitted parameters fixed, the number of families with an estimated *P* < 0.05 ranged from 192 to 837. Because our tree has four taxa in two pairs below the root, we can sum the distribution exactly. We therefore computed each *P* value exactly under the fitted model, as CAFE 5 defines it, and considered families with *P* < 0.05 significant.

To test whether the CAFE results reflected losses during annotation transfer, we binned orthogroups by their number of copies in the reference. For each bin, we computed the mean share of reference copies retained in the three transferred ingroup genomes and the share of significant families, and compared families with one reference copy with those with two or more using a two-sided Fisher’s exact test. Among families with at least one reference copy, we used a two-sided Mann-Whitney test to compare the copy deficit of significant families with that of the others. We also correlated the net number of contractions on each lineage with that lineage’s total copy deficit relative to the reference.

### Synteny

We derived synteny from the gene coordinates of the annotations described above. For each pair of ingroup assemblies, anchors were genes, coding or non-coding, that shared an identifier on the three chromosome-scale sequences of both. We paired chromosomes by the number of shared anchors and named them chr1 to chr3 after the *quinquefasciatus* chromosomes. Assemblies deposit chromosomes in either orientation, so we oriented each pair so the longer of its increasing and decreasing anchor runs was increasing. We defined collinearity as the share of anchors on the longest increasing run. We defined an inversion as a run of at least three anchors in decreasing order spanning at least 100 kb. We joined inversions from different pairs into recurrence clusters by single linkage when their anchor gene sets had a Jaccard index of at least 0.5. We counted anchors on nonhomologous chromosomes separately and excluded them from the collinearity and inversion counts.

### Repeats and Key Gene Families

We built a repeat library from the *quinquefasciatus* assembly with RepeatModeler 2.0.7 (Flynn et al. 2020) and used it to mask the four ingroup assemblies with RepeatMasker 4.1.7 (Smit et al. 2013–2015). We annotated proteins with eggNOG mapper 2.1.12 (Cantalapiedra et al. 2021) against the eggNOG 5.0 database (Huerta-Cepas et al. 2019). Nine families were then picked out from the eggNOG descriptions and gene names by weighted keyword patterns: cytochrome P450s, glutathione S transferases, carboxylesterases, odorant, gustatory and ionotropic receptors, odorant binding and chemosensory proteins, and immune genes of the classes cataloged by Bartholomay et al. (2010) (Toll-like receptors, peptidoglycan recognition proteins, thioester-containing proteins, IMD and antimicrobial peptides). We assigned an orthogroup to the family with the most annotated members, provided that family accounted for at least two genes or a quarter of the orthogroup’s ingroup genes. Counts are the numbers of genes per form in the assigned orthogroups.

### Anvi’o Pangenome

To check the OrthoFinder partition with a second clustering method, we clustered the same 57,815 ingroup proteins with Anvi’o 9 (Eren et al. 2021), following Delmont and Eren (2018). Each gene became a contig containing its coding sequence, with an external gene call that carried the protein used by OrthoFinder, so Anvi’o clustered those proteins rather than its own translations. It compared all of them with DIAMOND 2.2.8 (Buchfink et al. 2021) and removed weak matches using the minbit criterion: the bit score of a match divided by the smaller of the two self scores, with a cutoff of 0.5 (Benedict et al. 2014). It then clustered the proteins with MCL (van Dongen and Abreu-Goodger 2012) at an inflation of 10, the value the anvi’o developers use for very closely related genomes, and aligned each gene cluster with FAMSA 2.2.2 (Deorowicz et al. 2016). We classed gene clusters by the number of forms they contained (all four, two or three, or one) and traced each gene to its orthogroup, its reference gene, and its Liftoff ORF flag. The commands are in workflow/scripts/anvio_pangenome.sh, and the comparison is in workflow/scripts/anvio_compare.py.

### Workflow and Reproducibility

All analyses except the anvi’o pangenome were run with a Snakemake workflow (Mölder et al. 2021), available at https://github.com/tylermaire/cx_pipiens_pangenome (release v3.0). The workflow’s patch script computed the exact CAFE *P* values from the stored CAFE output by running the same code as the workflow rule. The supplementary tables and a table in the repository (results/manuscript_values.tsv) provide each reported value with its source file, and the run notes in the repository (docs/v5_run.md) record the earlier run and the checks performed outside the workflow. Claude (Anthropic), an AI assistant, was used to help write and review the workflow code; the authors checked all code and analyses.

## Results

### Assemblies and Transferred Gene Sets

The four ingroup assemblies ranged from 533 to 573 Mb, and the *Cx. perexiguus* assembly was 436 Mb (Table 1). Contiguity varied far more. Contig N50 ranged from 0.37 Mb in *molestus* to 3.26 Mb in *pipiens*. In genome mode, BUSCO completeness was 95.2 to 98.8% for the ingroup and 98.4% for *Cx. perexiguus*, and duplicated BUSCO genes were more frequent in the *pallens* and *molestus* assemblies (4.5 and 4.4%) than in the *quinquefasciatus* and *pipiens* assemblies (1.0 and 0.4%) (Supp. Table S1).

The three transferred ingroup gene sets were smaller than the reference, with 14,106 to 14,489 protein-coding genes against 15,094 in *quinquefasciatus* (Table 2). Protein-mode completeness shows the cost of transfer: it dropped 14.6 to 17.8 points below genome-mode completeness in the transferred ingroup genomes (to 77.4 to 81.6%), but only 0.7 points in the reference (95.2%) and 3.8 points in the independently annotated *Cx. perexiguus* gene set (94.6%).

**Table 2.** Gene sets after annotation transfer and their place in the pangenome.

| Form | Protein coding genes | Complete BUSCO, protein (%) | Without a valid ORF (%) | Without a valid ORF, clean reference models (%) | Core | Shell | Cloud | Unassigned |
| --- | --- | --- | --- | --- | --- | --- | --- | --- |
| *Cx. quinquefasciatus* (reference) | 15,094 | 95.2 | – | – | 12,279 | 2,492 | 221 | 102 |
| *Cx. pipiens pallens* | 14,106 | 80.6 | 30.1 | 26.7 | 11,602 | 2,029 | 25 | 450 |
| *Cx. pipiens* f. *molestus* | 14,126 | 77.4 | 34.3 | 31.1 | 11,610 | 1,867 | 23 | 626 |
| *Cx. pipiens* f. *pipiens* | 14,489 | 81.6 | 32.9 | 29.6 | 11,679 | 2,282 | 34 | 494 |
| *Cx. perexiguus* | 12,534 | 94.6 | – | – | 9,044 | 1,979 | 301 | 998<sup>a</sup> |

One protein per gene (longest isoform). Valid open reading frames (ORFs) as recorded by Liftoff; the last ORF column sets aside models whose *Cx. quinquefasciatus* source model is partial or carries a RefSeq sequence exception (872 of the kept reference models). *Culex perexiguus* keeps its own Ensembl gene set, which carries no Liftoff flags. Core, shell, and cloud show the number of genes of each form in orthogroups within each compartment, classified by the four ingroup forms; unassigned genes were placed in no orthogroup.
<sup>a</sup> further 212 *Cx. perexiguus* genes are in the 67 outgroup-only orthogroups.

About a third of the transferred models were broken. Liftoff flagged 30.1% of the kept models in *pallens*, 34.3% in *molestus,* and 32.9% in *pipiens* as lacking a valid ORF, and set aside 26.7 to 31.1% of models with partial or discrepant reference models (Table 2, Supp. Table S2). The most common defect was a missing stop codon (17.6 to 19.2% of models). Missing start codons came next (10.4 to 12.9%), and internal stop codons were rare (2.0 to 2.3%).

### Pangenome Partition and the Composition of the Cloud

OrthoFinder placed 56,143 of the 57,815 ingroup genes (97.1%) in orthogroups. Of the 13,672 orthogroups, 13,605 contained ingroup genes: 10,228 (75.2%) were core, 3,113 (22.9%) shell, and 264 (1.9%) cloud, and the other 67 held only *Cx. perexiguus* genes (Fig. 1a, b; Supp. Table S3). Cloud orthogroups numbered 201 in *quinquefasciatus* but only 28 in *pipiens*, 18 in *molestus,* and 17 in *pallens*.

**Fig. 1.**
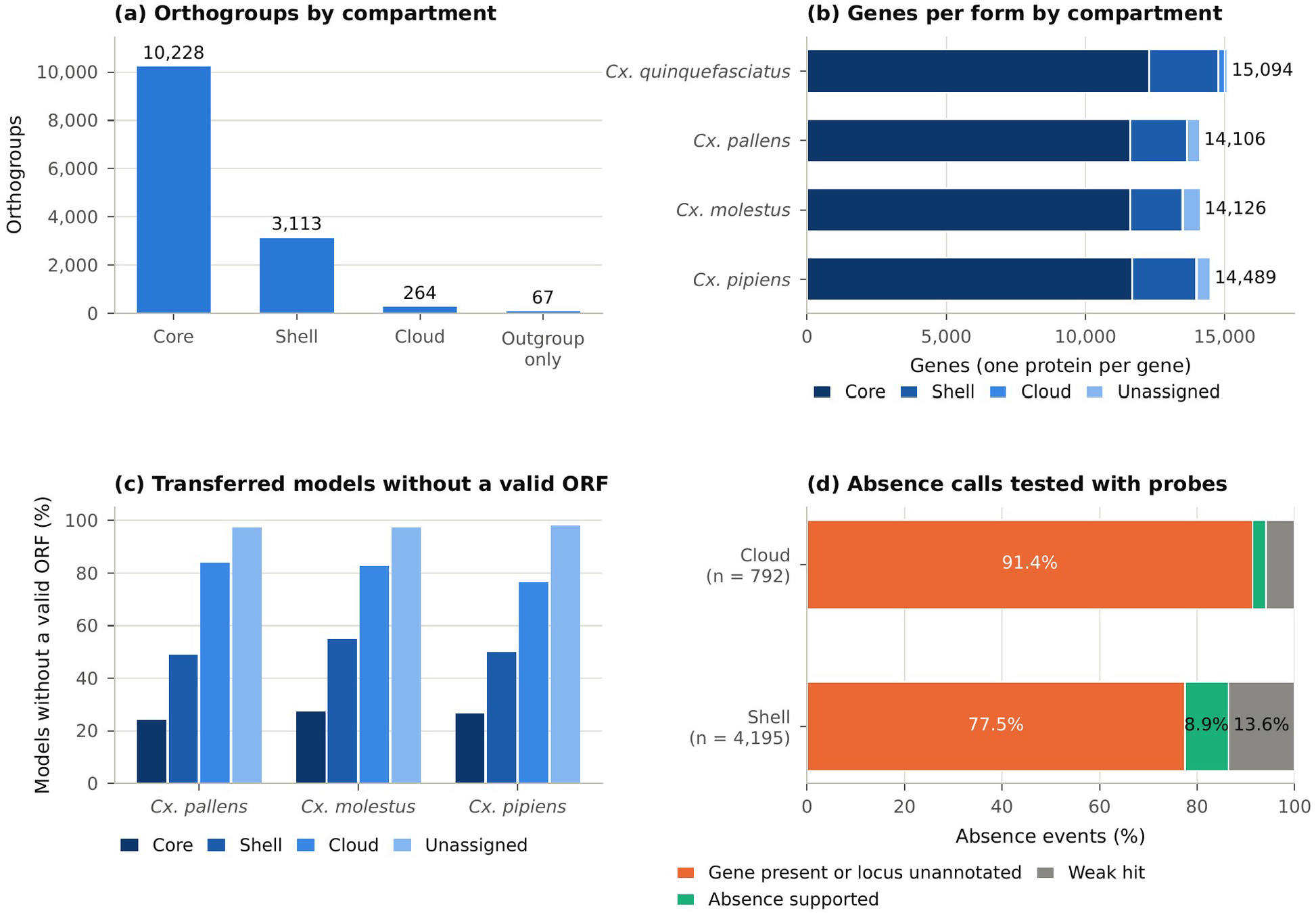
Pangenome partition of the four *Cx. pipiens* complex forms, and tests of the cloud. (a) Orthogroups by compartment over the four ingroup forms: core (genes from all four forms), shell (two or three), cloud (one), and outgroup only (*Cx. perexiguus* genes only). (b) Genes of each form (one protein per gene) in each compartment, plus the genes that OrthoFinder assigned to no orthogroup. (c) Share of the kept transferred models that Liftoff flagged as lacking a valid open reading frame, by compartment, in the three ingroup genomes annotated by transfer. (d) Outcome of testing every absence implied by cloud and shell orthogroups, by gene identifier and by protein (miniprot) and genomic (minimap2) alignment. An artifact is a model of the same gene present in another orthogroup or in none, or the locus found without a usable model. A supported absence has no hit, or hits only on other genes. Weak hits fell below the thresholds.

The cloud is not a set of genes unique to a form (Supp. Table S4). Each transferred model is a copy of a reference gene, so none of the 63 cloud orthogroups in the three transferred ingroup genomes can contain a gene missing from *quinquefasciatus*. Each exists because a transferred model failed to cluster with its source model.

The 201 cloud orthogroups of *quinquefasciatus* need more explanation. In 186 of them, the same gene had transferred models in other forms, clustered elsewhere, and only 15 (7.5%) had no counterpart. Usually, the reference model clustered with its *Cx. perexiguus* ortholog, while its transferred copies did not. A *Cx. perexiguus* gene sat in 197 of these 201 orthogroups, and in 244 of all 264 cloud orthogroups (92.4%). Most cloud orthogroups (233) contained a single ingroup gene, and OrthoFinder places a single gene in an orthogroup only if it is the only gene from another species.

The share of models without a valid ORF rose from 24.1 to 27.5% in core orthogroups to 48.8 to 54.8% in shell, 76.5 to 84.0% in cloud, and 97.3 to 98.0% among unassigned genes (Fig. 1c, Supp. Table S2). Most cloud genes are broken transfers, or reference genes whose transfers broke, rather than genes unique to a form.

Testing the absences directly gave the same answer (Fig. 1d, Supp. Table S5). Cloud orthogroups implied 792 absence events, and 91.4% of them were artifacts. In 701 events, the target genome held a model of the same reference gene, placed in another orthogroup or, in 137 events, in none; in 23 more, the sequence was present at a locus with no usable model. The two models of the same gene often differed in sequence (median protein identity 0.23), as expected from broken transfers. Only 21 events (2.7%) supported an absence (8 absent and 13 paralog only), and 47 (5.9%) were weak. Of the 4,195 events implied by shell orthogroups, 77.5% were artifacts, 8.9% supported an absence, and 13.6% were weak.

### Four Taxon Relationships and Gene Tree Discordance

Of 9,168 single-copy orthologs, 9,126 alignments survived trimming (4,928,665 amino acid columns, 1.09% gaps or ambiguous residues). The concatenated tree separated *pallens* and *quinquefasciatus* from *molestus* and *pipiens* with full UFBoot support. Many loci disagreed. Only 4,491 of 7,621 gene trees (58.9%) and 45.1% of 79,880 parsimony informative sites supported this split (Fig. 2b, Supp. Table S6). We could not build gene trees for 1,505 alignments, all of which had fewer than four distinct sequences (794 were invariant).

**Fig. 2.**
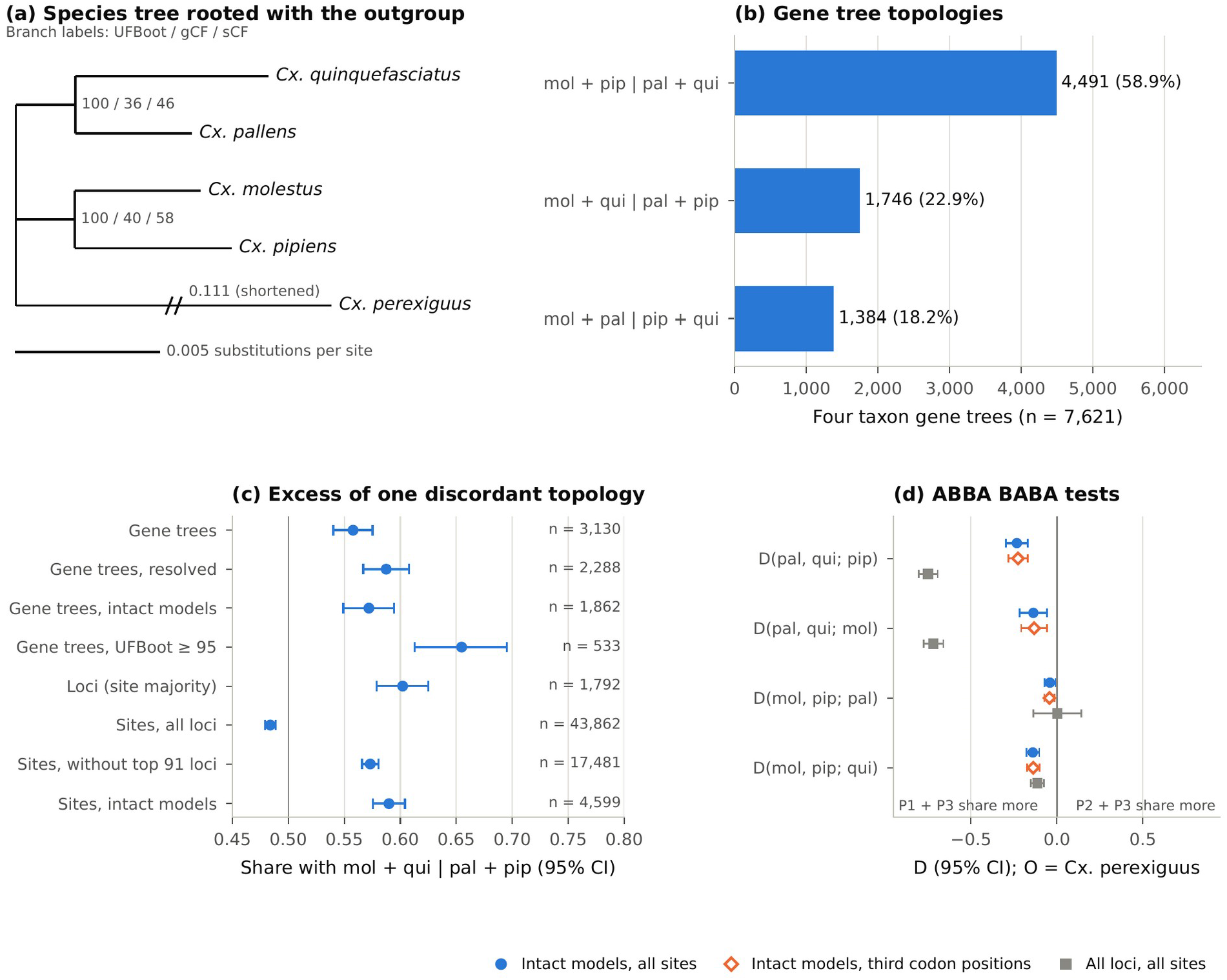
Relationships among the forms, gene tree discordance and ABBA-BABA tests. (a) Species tree rooted with *Cx. perexiguus*, from concatenated protein alignments of 4,592 five-taxon loci with four intact ingroup gene models. Branches are labeled with ultrafast bootstrap support (UFBoot), gene concordance factor (gCF), and site concordance factor (sCF); branch lengths are in amino acid substitutions per site, and the outgroup branch is shortened. (b) Number of four-taxon gene trees supporting each of the three unrooted topologies (mol, *molestus*; pip, *pipiens*; pal, *pallens*; qui, *quinquefasciatus*). (c) Share supporting mol + qui | pal + pip, with exact 95% confidence intervals and the numbers of discordant trees, loci, or sites. The share is shown among discordant gene trees (all gene trees; gene trees whose internal branch is longer than IQ-TREE’s minimum, resolved; gene trees of loci whose four gene models are intact; gene trees with UFBoot of at least 95), among loci counted once each by the split with the most parsimony informative sites, and among parsimony informative sites summed over all loci, over all but the 91 most variable loci (1% of loci) and over loci with intact models. Under incomplete lineage sorting alone, it is expected to be 0.5. (d) D statistics D(P1, P2; P3) with *Cx. perexiguus* as the outgroup and 95% confidence intervals from a block jackknife, on loci with intact models (all sites and third codon positions) and on all loci (all sites). D < 0 means that P1 and P3 share more derived alleles than P2 and P3.

The two discordant topologies were not equally frequent (Fig. 2b). Gene trees pairing *molestus* with *quinquefasciatus* and *pallens* with *pipiens* (1,746, 22.9%) outnumbered those pairing *molestus* with *pallens* and *pipiens* with *quinquefasciatus* (1,384, 18.2%), making up 55.8% of discordant trees (95% CI 54.0 to 57.5%; binomial *P* = 1.0 × 10^−10^). The excess grew with gene tree support, reaching 65.5% among gene trees with UFBoot of at least 95 (349 against 184; 95% CI 61.3 to 69.5%; *P* = 8.0 × 10^−13^) (Fig. 2c).

Many gene trees were unresolved. In 1,277 (16.8%) trees, the internal branch had the minimum length, and these splits were almost evenly distributed among the three topologies (435, 402, and 440). Among the 6,344 resolved gene trees, 63.9% supported the species tree, and the excess topology made up 58.7% of discordant trees (1,344 against 944; *P* = 6.1 × 10^−17^). Loci with four intact gene models also showed the excess, at 57.2% of their 1,862 discordant gene trees (*P* = 5.7 × 10^−10^) and 77.8% of those with UFBoot of at least 95 (151 against 43; *P* = 2.8 × 10^−15^). One subset lacked it. The 1,065 gene trees with a terminal branch longer than 0.1 split 289 against 273 (*P* = 0.53), whereas the other 6,556 gene trees gave 56.7% (1,457 against 1,111) (Supp. Table S6).

Summed over all loci, parsimony-informative sites leaned the other way (21,224 against 22,638; 48.4%; *P* = 1.5 × 10^−11^). But informative sites were concentrated in a few loci. The median locus had two informative sites, while the 91 most variable loci (1% of loci) held 45.6% of all informative sites. In those loci, the species-tree split accounted for only 27.6% of informative sites, less than the minor discordant split (41.7%), and every one contained an incomplete gene model, compared with 34.0% of other loci. When each locus was counted once, by the split with the most informative sites, 1,079 loci supported the excess topology and 713 the alternative (60.2%; 95% CI 57.9 to 62.5%; *P* = 5.1 × 10^−18^), and without the 91 most variable loci, 57.3% of 17,481 discordant informative sites supported it (*P* = 1.1 × 10^−83^). In the 5,959 loci with intact models, 74.4% of informative sites supported the species tree split, and the excess topology held 59.0% of discordant sites (2,713 against 1,886; *P* = 2.7 × 10^−34^; Fig. 2c) and 62.5% of locus votes (610 against 366; *P* = 5.3 × 10^−15^; Supp. Table S6).

### The Root and ABBA-BABA Tests

Of the 6,831 five-taxon loci with trimmed alignments of at least 50 columns, 4,592 had four intact ingroup models. Rooted with *Cx. perexiguus*, the tree from these loci (2,444,417 amino acid columns) placed the root between (*molestus*, *pipiens*) and (*pallens*, *quinquefasciatus*), with full UFBoot support for both pairs (Fig. 2a, Supp. Table S12). Concordance was modest: gCF was 40.18 and sCF 58.26 for (*molestus*, *pipiens*), and gCF 35.50 and sCF 46.05 for (*pallens*, *quinquefasciatus*). Still, the species tree was the most frequent of the 15 rooted topologies among the 4,017 gene trees of these loci (21.8%; loci with too few distinct sequences gave no tree), and among the 2,150 with fully resolved internal branches (26.2%). Among resolved gene trees, the next most frequent grouping was *pallens* with (*molestus*, *pipiens*) (17.7%). Overall, in gene trees, *quinquefasciatus* diverged first in 1,215 (30.2%) and *pallens* in 574 (14.3%).

Adding loci with incomplete models moved the root. The tree from all 6,831 loci placed it on the *quinquefasciatus* branch, grouping *pallens* with (*molestus*, *pipiens*), again with full UFBoot support (Supp. Table S13). The site counts show why. Sites where *pallens*, *molestus,* and *pipiens* shared a derived state against *quinquefasciatus* and the outgroup numbered 256 per locus in the 408 loci whose reference model carried a RefSeq exception, and 53 per locus in the 381 loci where all three transferred models lacked a valid ORF, against 4 per locus in intact loci. Where RefSeq had corrected the reference model, the three transferred models typically carried the same frameshift relative to the reference model and the outgroup. Their misaligned codons then appeared as sites where the three forms shared a derived state. Errors shared by two forms inflated that pair similarly. In loci where only the *pallens* and *pipiens* models lacked a valid ORF, sites with a derived state in those two forms alone averaged 66 per locus, against 1.3 in intact loci.

On loci with intact models, *pallens* shared more derived alleles with *pipiens* than *quinquefasciatus* did (D(*pallens*, *quinquefasciatus*; *pipiens*) = −0.233, Z = −7.2, corrected *P* = 1.9 × 10^−12^). It also shared more with *molestus*, less strongly (D(*pallens*, *quinquefasciatus*; *molestus*) = −0.137, Z = −3.3, *P* = 0.003) (Fig. 2d, Supp. Table S12). Form *molestus* shared more derived alleles with *quinquefasciatus* than *pipiens* did (D(*molestus*, *pipiens*; *quinquefasciatus*) = −0.140, Z = −7.3, *P* = 9.3 × 10^−13^), whereas *molestus* and *pipiens* differed little in the alleles they shared with *pallens* (D(*molestus*, *pipiens*; *pallens*) = −0.041, Z = −2.4, *P* = 0.062).

Third codon positions gave nearly the same numbers (D = −0.226, −0.132, −0.138 and −0.043, in the order above), although there the slight excess shared by *molestus* and *pallens* in the last test was significant (corrected *P* = 0.021). Loci whose four ingroup sequences were at least 95 or 98% identical gave values of the same sign, similar or larger for three of the tests but mostly smaller for D(*molestus*, *pipiens*; *pallens*) (−0.018 to −0.042, against −0.041), and at 98% identity D(*pallens*, *quinquefasciatus*; *molestus*) was no longer significant (Z = −2.4 and −2.3) (Supp. Tables S12 and S13).

On all 6,846 loci, shared transfer errors pushed D(*pallens*, *quinquefasciatus*; *pipiens*) and D(*pallens*, *quinquefasciatus*; *molestus*) to −0.749 and −0.719. D(*molestus*, *pipiens*; *quinquefasciatus*) changed less (D = −0.113), as expected, because in that test an error shared by two or all three transferred forms produces neither an ABBA nor a BABA site.

### Divergence Between Forms

The concatenated four-taxon tree had terminal branches of 0.029 (*pallens*) to 0.054 (*quinquefasciatus*) substitutions per site, and an internal branch of 0.012. The per-locus gene trees were much shorter. Their median terminal branches were 7 to 9 times shorter (0.0032 to 0.0061), and the median internal branch was about 2.4 times shorter (0.005). For each form, 3.3 to 5.9% of gene trees had a terminal branch longer than 0.1, and 16.9 to 26.4% had a terminal branch at IQ-TREE’s minimum length, with no substitution inferred. Median pairwise protein identity across the single-copy alignments was 98.5 to 99.1% (Table 3, Supp. Table S7), although 12.3 to 19.2% of loci fell below 95%.

**Table 3.** Pairwise identity and synteny between the ingroup assemblies.

| Pair | skani ANI (%) | skani aligned fraction (%) | minimap2 identity (%) | Median protein identity (%) | Anchors on homologous chromosomes | Collinear anchors (%) | Shared genes on another chromosome (%) | Inversions $\geq 100$ kb |
| --- | --- | --- | --- | --- | --- | --- | --- | --- |
| *quinquefasciatus* vs *pallens* | 94.22 | 24.2–24.5 | 94.33 | 99.10 | 14,210 | 85.4 | 12.1 | 77 |
| *quinquefasciatus* vs *molestus* | 93.19 | 18.9–19.4 | 92.88 | 98.69 | 13,762 | 83.8 | 13.4 | 112 |
| *quinquefasciatus* vs *pipiens* | 92.80 | 17.4–18.7 | 93.28 | 98.52 | 15,259 | 88.1 | 10.0 | 49 |
| *pallens* vs *molestus* | 93.55 | 20.6–20.9 | 93.26 | 98.88 | 12,809 | 83.1 | 13.2 | 129 |
| *pallens* vs *pipiens* | 93.33 | 20.6–21.9 | 93.79 | 98.81 | 13,879 | 87.9 | 11.5 | 59 |
| *molestus* vs *pipiens* | 94.35 | 27.4–28.8 | 94.66 | 99.10 | 13,554 | 85.4 | 12.6 | 109 |

skani ANI and aligned fraction (both directions) on the complete assemblies; minimap2 identity is the length-weighted identity of alignments between the three chromosome-scale sequences; median protein identity is over the 9,126 trimmed single-copy ortholog alignments (9,114 to 9,126 per pair, those with columns where both forms have a residue). Anchors are genes, coding or non-coding, shared by both assemblies on homologous chromosomes. Further values are in Supp. Tables S7 and S9.

Whole-genome measures came out lower. skani ANI ranged from 92.80% (*pipiens* against *quinquefasciatus*) to 94.35% (*molestus* against *pipiens*). Still, it was computed over only 17 to 29% of each assembly, and minimap2 alignment identity over the chromosome-scale sequences ranged from 92.9 to 94.7% (Table 3). For pairs including *Cx. perexiguus*, skani returned no estimate. All three measures ranked the two sister pairs as the most similar. They disagreed on the least similar pair: skani ANI and protein identity ranked *pipiens* against *quinquefasciatus* lowest, while minimap2 identity ranked *molestus* against *quinquefasciatus* lowest.

### Gene Family Size and the Transfer Control

CAFE estimated a birth and death rate of 0.045 and a gamma shape of 0.271. Of the 12,864 families it tested, 264 had *P* < 0.05 (Supp. Table S8). On the terminal branches, decreases dominated in the transferred genomes: 61 increases against 1,049 decreases in *pallens*, 131 against 1,176 in *molestus,* and 158 against 746 in *pipiens*, compared with 473 and 582 in *quinquefasciatus*. Both internal branches showed only increases (53 and 106).

These changes follow copy loss during transfer (Fig. 3). In families with one reference copy, the transferred ingroup genomes retained a mean share of 0.94 of the reference copies. That share fell to 0.78, 0.74, 0.71, and 0.71 for families with 2, 3 to 5, 6 to 10, and more than 10 reference copies, and over the same bins the share of significant families rose from 0.11% to 11.2, 22.8, 33.3, and 68.2%. Families with two or more reference copies were significantly more common than families with one (182 of 1,108 against 12 of 11,188; odds ratio 183, relative risk 153; *P* = 5.3 × 10^−179^). Significant families carried a larger mean copy deficit (1.95 against 0.10 copies; *P* = 3.8 × 10^−175^), and the net contraction of each lineage tracked its total copy deficit (Pearson *r* = 0.993, *n* = 4). Families with no reference copy: 568 tested orthogroups that, by construction, contain only transferred models separated from their source genes contributed 70 significant families (12.3%).

**Fig. 3.**
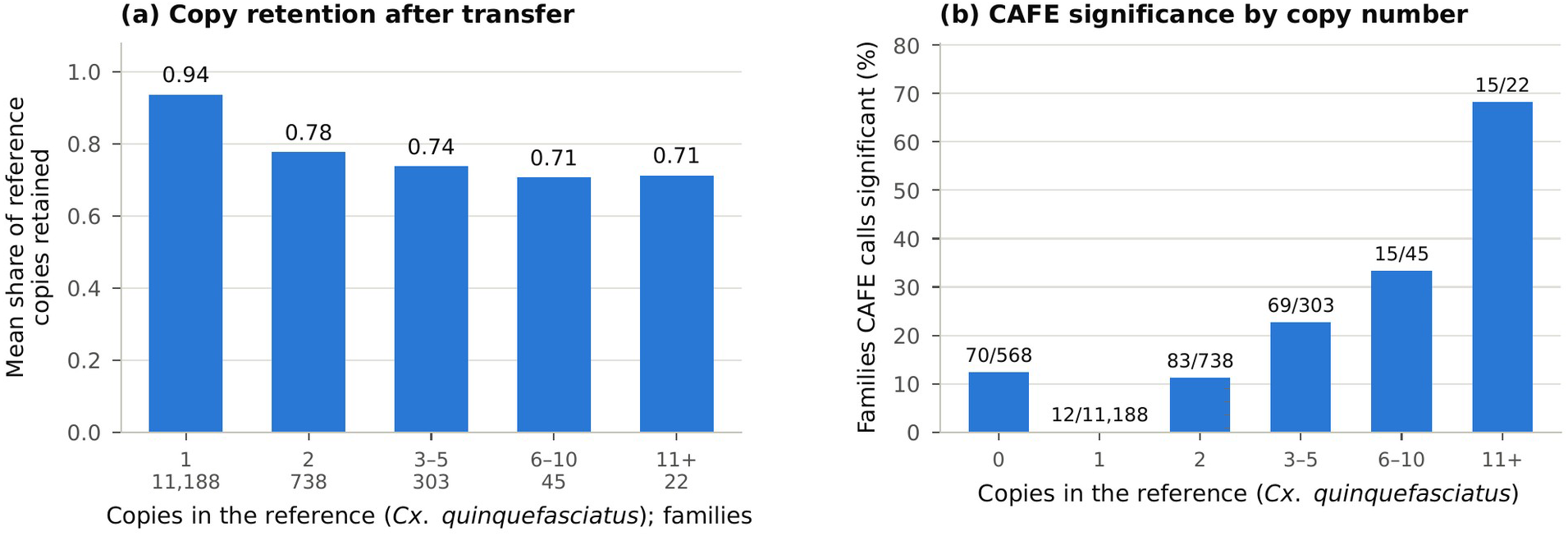
Gene family size and annotation transfer. CAFE groups the 12,864 orthogroups by the number of copies in the *Cx. quinquefasciatus* reference. (a) Mean share of reference copies retained in the three ingroup genomes annotated by transfer, for families with at least one reference copy, with the number of families in each group below the axis. (b) Share of families with *P* < 0.05 (exact *P* values under the fitted CAFE model), with the numbers of significant and total families.

In this run, CAFE’s own simulated *P* values would have added 572 more families, each with one copy in one form of each pair and none in the other two, all with an exact *P* of 0.055 (Supp. Table S8). For these reasons, we do not read the CAFE results as gene family evolution.

### Synteny

The three chromosomes corresponded one-to-one in every pair of ingroup assemblies, and in 12 of the 18 chromosome pairs one assembly was deposited in the opposite orientation (Fig. 4, Supp. Table S9). Between 12,809 and 15,259 anchors per pair lay on homologous chromosomes, and 83.1-88.1% were collinear. Another 10.0 to 13.4% of shared genes lay on a non-homologous chromosome in one of the two assemblies. These were scattered across all chromosomes, with many near the middle of each chromosome (Fig. 4).

**Fig. 4.**
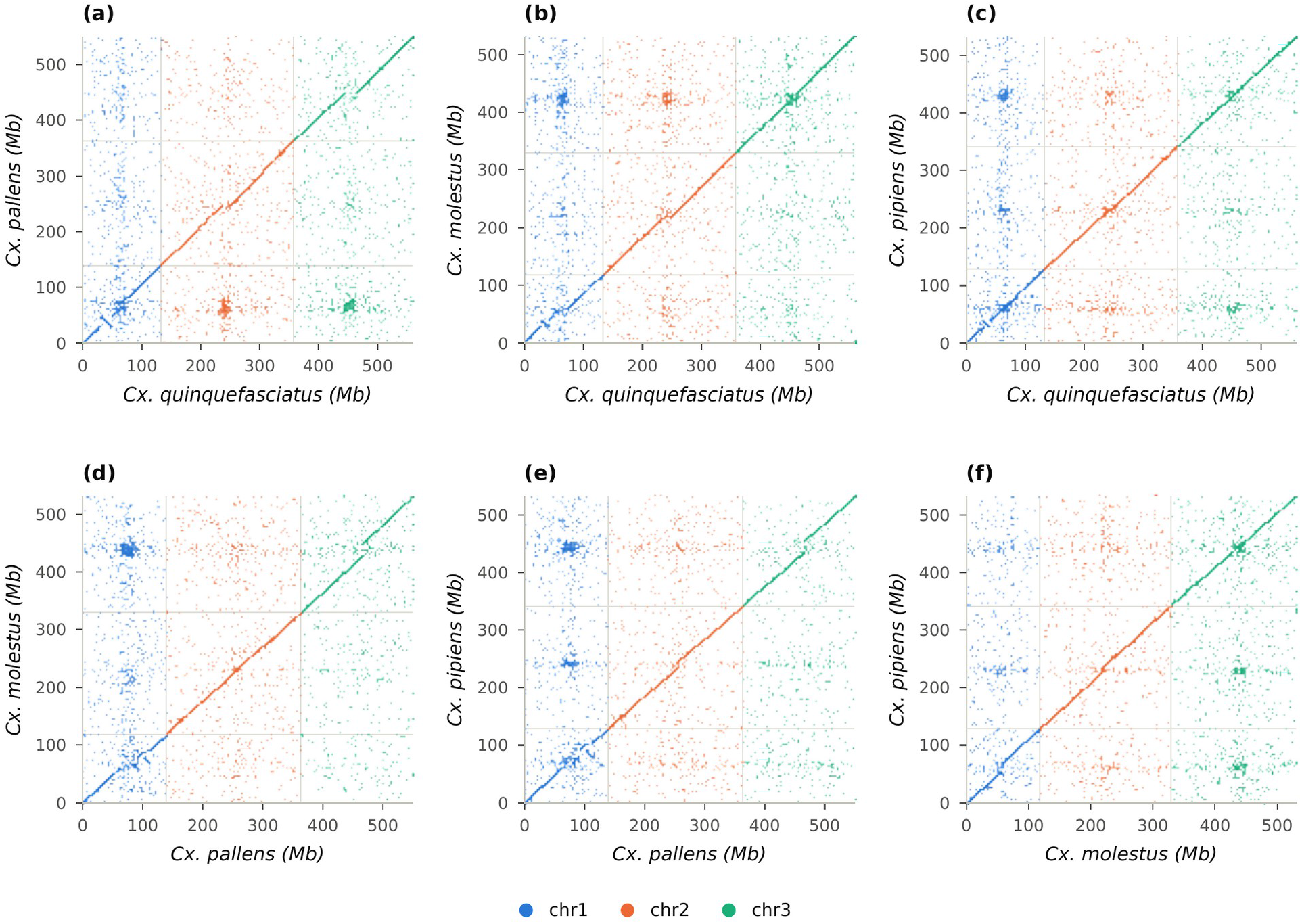
Pairwise synteny of the four ingroup assemblies based on shared-gene positions. Each point is a gene, coding or non-coding, shared by two assemblies, plotted at its positions on the three chromosome-scale sequences of each, concatenated in chromosome order (chr1 to chr3, numbered after *Cx. quinquefasciatus*). We reversed chromosomes deposited in the opposite orientation in the assembly on the y-axis. Points are colored by assembly chromosome (x-axis); points off the diagonal blocks are genes placed on a non-homologous chromosome in one of the two assemblies. (a to c) *Cx. quinquefasciatus* against *pallens*, *molestus,* and *pipiens*; (d, e) *pallens* against *molestus* and *pipiens*; (f) *molestus* against *pipiens*.

We found 535 inversions of at least 100 kb, from 49 (*quinquefasciatus* against *pipiens*) to 129 (*pallens* against *molestus*) per pair. Of these, 23 spanned at least 1 Mb, the largest 1.81 Mb on chr1 between *quinquefasciatus* and *pallens*. The inversions formed 391 recurrence clusters, and 95 of the clusters appeared in two or three pairs. Forty-seven appeared in all three pairs that include one assembly, the pattern expected for a rearrangement in, or an assembly error of, that assembly: *molestus* in 31, *pallens* in 11, *quinquefasciatus* in 5, and *pipiens* in none. The other recurrent clusters appeared in two pairs (46) or in three pairs without a shared assembly (2). Nine recurrent clusters spanned at least 1 Mb.

### Repeats

Interspersed repeats covered 53.0-54.7% of the ingroup assemblies. With simple repeats (3.6 to 4.2%) and low-complexity sequence (0.15 to 0.18%), the total masked fraction was 57.4 to 59.0% (Table 1, Supp. Table S10). For the *quinquefasciatus* assembly, the masked fraction was 58.0%, close to the 58.7% reported for this assembly by Ryazansky et al. (2024) and about twice the 29% reported for the earlier CpipJ2 assembly (Arensburger et al. 2010).

### Anvi’o Pangenome

Anvi’o placed 57,716 of the 57,815 ingroup proteins in 16,623 gene clusters (Fig. 5, Supp. Table S14). The other 99, mostly fragments (median length 12 residues), had no DIAMOND hit, not even to themselves, so they never entered the clustering; OrthoFinder also left 97 of them unassigned. Of the gene clusters, 9,465 (56.9%) contained genes from all four forms, 4,142 from two or three forms, and 3,016 from one (Fig. 5a), and 8,577 held exactly one gene of each form.

**Fig. 5.**
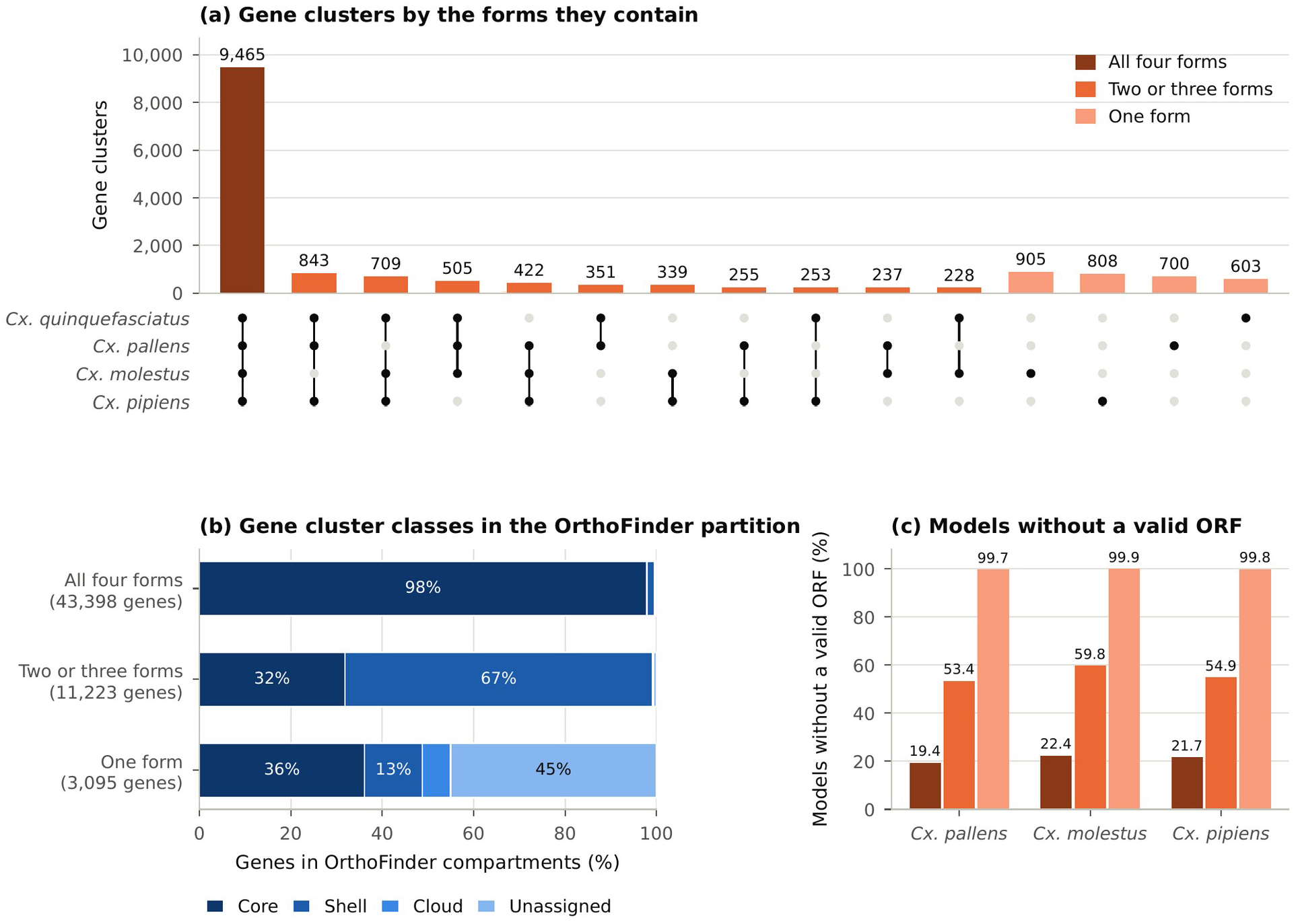
Anvi’o gene clusters of the four ingroup forms, from the 57,815 proteins of the OrthoFinder partition. (a) Gene clusters by the combination of forms whose genes they contain, shown by the matrix below the bars, and shaded by class: all four forms, two or three forms, or one form. (b) Genes of each class by OrthoFinder compartment: core, shell, and cloud orthogroups, and genes that OrthoFinder left unassigned. (c) Share of transferred models without a valid ORF (Liftoff) in each class, for the three ingroup genomes annotated by transfer, shaded as in (a). The 99 proteins without a DIAMOND hit, which anvi’o excludes from gene clusters, are not included.

On the core, the two methods agreed. Of the 43,398 genes in four-form clusters, 97.9% were in core orthogroups, and 9,086 of the 10,228 core orthogroups fell entirely in four-form clusters. Overall, 89.4% of clustered genes landed in the matching OrthoFinder compartment (core for clusters of all four forms, shell for two or three, and cloud or unassigned for one; Fig. 5b).

The methods differed in how they grouped broken models. Of the 13,851 transferred models without a valid ORF, 5,703 (41.2%) fell outside the gene cluster of their reference gene’s model and 3,794 (27.4%) outside its orthogroup; for models with a valid ORF, the shares were 0.2 and 0.4%. Anvi’o, in other words, separated broken models from their source genes more often than OrthoFinder did. The 3,016 one-form clusters (603 in *quinquefasciatus*, 700 in *pallens*, 905 in *molestus* and 808 in *pipiens*) contained 3,095 genes, of which OrthoFinder left 45.1% unassigned and placed 36.1% in core orthogroups. In the transferred genomes, nearly all of these genes were broken. Of 2,445 such genes, 2,440 (99.8%) lacked a valid ORF, compared with 19.4 to 22.4% of transferred models in four-form clusters and 53.4 to 59.8% in clusters of two or three forms (Fig. 5c). The reference models in one-form clusters cannot be broken transfers, so we looked at their genes in the other forms instead. Of the 650 *quinquefasciatus* genes in one-form clusters, 585 had kept models in other forms; 582 included a model without a valid ORF, and 200 (30.8%) had RefSeq models that were partial or carried an exception, compared with 5.8% of all reference models.

Overall, 3,022 of the 3,095 genes in one-form clusters (97.6%) were broken transfers or reference genes with a broken transfer. Under either method, the genes found in one form are mostly broken transfers or reference genes with broken transfers, rather than genes unique to a form. Both analyses rest on the same transferred models, so their agreement is not an independent test of gene content.

### Key Gene Families

Copy numbers in the nine families examined were similar among forms (Table 4, Supp. Table S11). *Cx. quinquefasciatus* had the highest or joint-highest count in every family; for example, 149 cytochrome P450s against 128 to 138 in the transferred genomes, and 120 odorant receptors against 103 to 104. Copy loss during transfer would produce exactly this pattern. These counts inventory the transferred gene sets and provide lower bounds for the transferred genomes, rather than evidence that the forms differ.

**Table 4.**
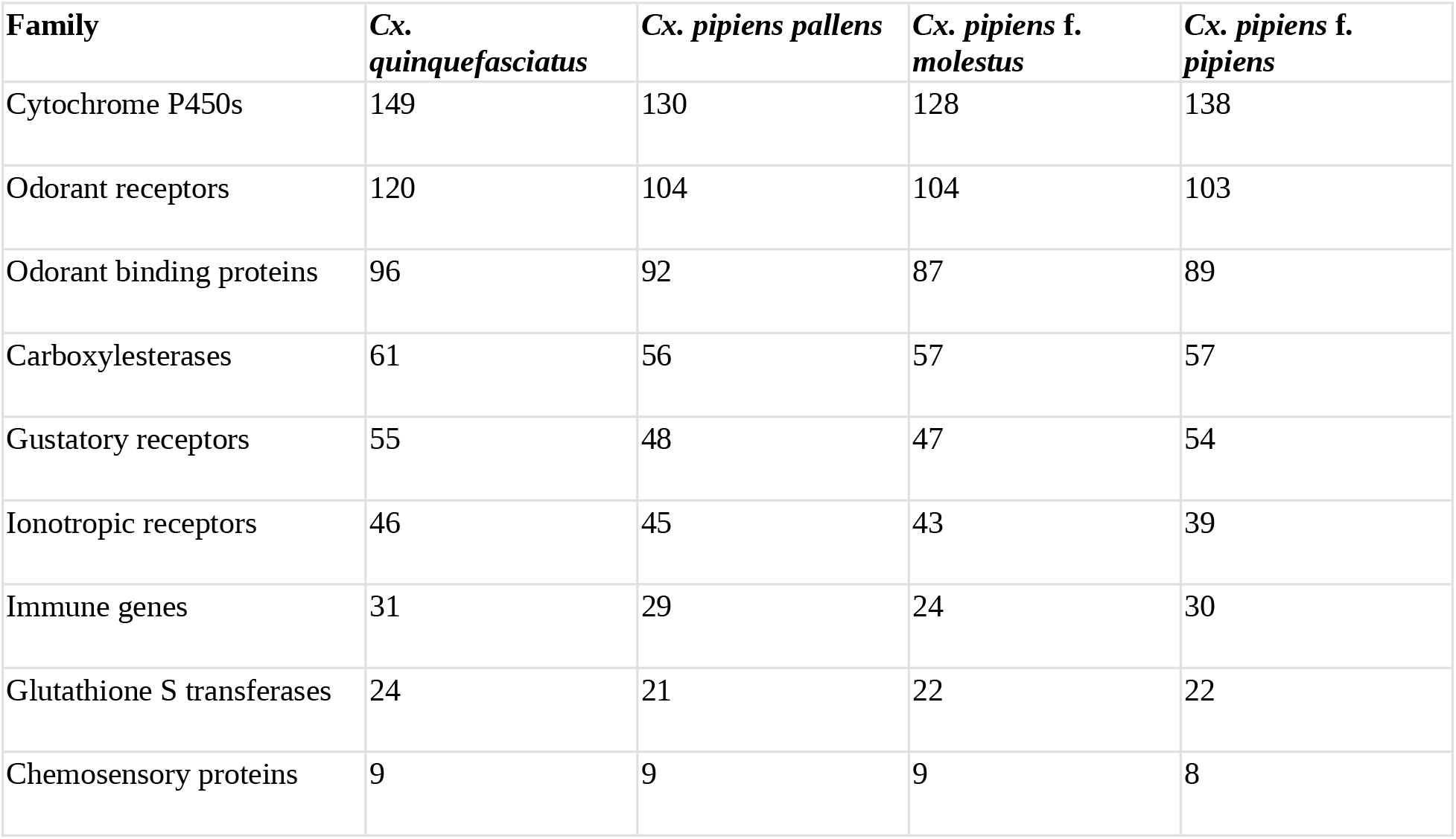
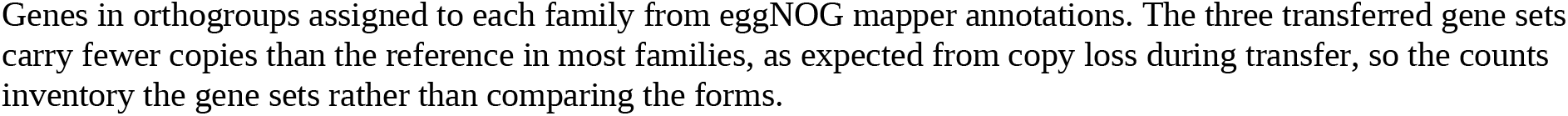
Genes assigned to key gene families in each form.

| Family | *Cx. quinquefasciatus* | *Cx. pipiens pallens* | *Cx. pipiens f. molestus* | *Cx. pipiens f. pipiens* |
| --- | --- | --- | --- | --- |
| Cytochrome P450s | 149 | 130 | 128 | 138 |
| Odorant receptors | 120 | 104 | 104 | 103 |
| Odorant binding proteins | 96 | 92 | 87 | 89 |
| Carboxylesterases | 61 | 56 | 57 | 57 |
| Gustatory receptors | 55 | 48 | 47 | 54 |
| Ionotropic receptors | 46 | 45 | 43 | 39 |
| Immune genes | 31 | 29 | 24 | 30 |
| Glutathione S transferases | 24 | 21 | 22 | 22 |
| Chemosensory proteins | 9 | 9 | 9 | 8 |

Genes in orthogroups assigned to each family from eggNOG mapper annotations. The three transferred gene sets carry fewer copies than the reference in most families, as expected from copy loss during transfer, so the counts inventory the gene sets rather than comparing the forms.

## Discussion

### Relationships Among the Forms and Gene Flow

Rooted with *Cx. perexiguus*, the four forms fall into two pairs, (*molestus*, *pipiens*) and (*pallens*, *quinquefasciatus*). The pairing of *pallens* with *quinquefasciatus* fits the evidence of *Cx. quinquefasciatus* ancestry in *pallens* (Fonseca et al. 2009, Aardema et al. 2020), and the pairing of the two forms of *Cx. pipiens* fits their taxonomy. Support for both pairs was modest, though. Only 59% of four taxon gene trees and 45% of informative sites supported the split between them, and in the rooted analysis, gCF was 40.2 and 35.5. This low concordance is expected when lineages diverge in quick succession, because incomplete lineage sorting leaves many gene trees discordant with the species tree (Degnan and Rosenberg 2009).

The imbalance between the two discordant topologies is harder to explain that way, since incomplete lineage sorting predicts they should occur at equal frequencies (Hibbins and Hahn 2022). Gene trees pairing *molestus* with *quinquefasciatus* and *pallens* with *pipiens* were in excess. The excess grew as we removed poorly supported gene trees, and it held among resolved gene trees. In loci with four intact gene models, this showed up in the gene trees, the informative sites, and the locus votes alike. Random estimation error and broken transferred models are therefore unlikely sources.

Systematic error cannot be excluded entirely. The most likely source, long branch attraction, would favor the other discordant topology, the one that joins the two longest terminal branches of the concatenated tree (*quinquefasciatus* and *pipiens*). Among gene trees with a terminal branch longer than 0.1, where this error should be strongest, the excess was absent rather than enriched. The summed count of informative sites across all loci leaned the other way. That count was dominated by 91 loci holding almost half of all informative sites, in which the species tree split had little more than a quarter of the sites, and every one of those loci contained a gene model that was not intact. Site concordance measures the share of sites supporting a split, so it depends on how variable the contributing loci are. In contrast, gene concordance estimates the share of the genome with each history (Lanfear and Hahn 2024). Here, gene trees are the better measure.

On its own, the excess cannot show which forms exchanged genes. In an unrooted tree of the four forms, putting *pallens* with *pipiens* also puts *molestus* with *quinquefasciatus*, so gene flow in either pair would make that tree more common (Hibbins and Hahn 2022). The ABBA-BABA tests do not have this problem because they use the outgroup to identify derived alleles and then ask which forms share them.

Form *molestus* shared more derived alleles with *quinquefasciatus* than *pipiens* did, pointing to gene flow between *molestus* and *quinquefasciatus*; members of the complex hybridize where they meet (Cornel et al. 2003, Kothera et al. 2009). *Culex pipiens pallens* shared more derived alleles than *quinquefasciatus* with both forms of *Cx. pipiens*, and the two forms differed little in the alleles they shared with *pallens*. That fits *Cx. pipiens* ancestry in *pallens* coming from the lineage of both forms, not from form *pipiens* alone. The weaker signal with *molestus* is expected. Gene flow between *molestus* and *quinquefasciatus* adds derived alleles shared by those two, and these offset part of the excess shared by *pallens* and *molestus*. The rooted gene trees agree: *quinquefasciatus* diverged first, with *pallens* grouped with the two forms of *Cx. pipiens*, twice as often as *pallens* diverged first.

These results support the view that *pallens* arose from hybridization between *Cx. quinquefasciatus* and *Cx. pipiens* (Aardema et al. 2020), with *quinquefasciatus* as its closer relative. They do not show when the exchange happened or which way genes moved. Ancestral population structure can also produce excess allele sharing (Durand et al. 2011, Lanfear and Hahn 2024). Resequencing several individuals per form would make it possible to find which parts of the genome were exchanged, and when (Hibbins and Hahn 2022, Suvorov et al. 2022).

The two tests of alleles shared by *pallens* call for caution. They compare transferred-gene models against the reference, which makes them most exposed to errors shared across transferred models; on all loci, those errors inflated them severalfold and moved the root to the *quinquefasciatus* branch. On intact loci, they kept their sign. Restricting those loci further to ingroup sequences at least 98% identical did not shrink them (−0.276 and −0.140). Still, we cannot rule out residual errors shared by the three transferred forms, which would favor both a tree with *quinquefasciatus* diverging first and negative values in these two tests. The test of alleles shared by *molestus* and *quinquefasciatus* is not exposed this way, since an error shared by two or all three transferred forms produces neither an ABBA nor a BABA site. It gave similar values for all loci and for intact loci.

### How Divergent Are the Forms?

Judged locus by locus, the forms have diverged very little. Median protein identity between forms was 98.5 to 99.1%, and median terminal branches were 0.003 to 0.006 substitutions per site. The concatenated tree gave terminal branches 7 to 9 times longer, because a few percent of loci with very long branches dominate the concatenated estimate. Whole-genome ANI is still lower, at 92.80-94.35%. But skani aligned only 17 to 29% of each assembly, and in assemblies that are more than half repeats, whole-genome identity mixes orthologous sequence with repeat copies and assembly differences. Those values describe how alike the assemblies are, not the divergence between the forms.

The per-locus values are approximate too. Errors in the transferred models add apparent substitutions, while the Liftoff identity threshold and the restriction to single-copy orthologs leave out the most divergent genes. Comparisons with divergence in other vector complexes, such as the *An. gambiae* complex (Fontaine et al. 2015), should use per locus measures from independently annotated genomes.

### What Transferred Annotations Can and Cannot Show

We transferred one annotation to each genome to avoid inflating lineage-specific genes by mixing annotation methods (Weisman et al. 2022). It did not protect us from inflation caused by transfer errors. About a third of the transferred models lacked a valid ORF, broken models piled up in the shell and cloud, and more than 90% of cloud absences were the same gene clustered elsewhere or left without a usable model. A second clustering method gave the same picture: almost every transferred model in a single form anvi’o gene cluster lacked a valid ORF. So the size of the cloud, and its spread among forms, say more about how well the reference annotation transferred to each assembly, and how the transferred models clustered with the outgroup, than about unique genes.

CAFE inferred widespread gene loss on the branches leading to the transferred genomes, and its significant families and contractions were concentrated in the families and lineages that lost the most copies in transfer. Annotation errors inflate gene family counts similarly in draft genomes (Denton et al. 2014). There is also a statistical problem. When many families share the same counts, as here, CAFE 5’s simulated *P* values can shift hundreds of families across a threshold from one run to the next; with few taxa, exact *P* values avoid this.

Transfer errors reach phylogenomic analyses too. Transferred models that share an error (for example, where RefSeq corrected the reference model for an error in the reference genome) carry the same misaligned codons, and those codons count as shared derived alleles. With all loci included, they placed the root on the *quinquefasciatus* branch with full bootstrap support. They inflated two D statistics severalfold, while the four-taxon gene tree test, which counts each locus once, was little affected. Studies that transfer one annotation to several genomes should check whether their phylogenomic signals concentrate in loci with broken or corrected models.

Questions about gene family evolution and genes specific to one form in this complex will need independent, evidence-based annotation of each genome. Two existing gene sets provide a starting point. The native RefSeq annotation of the *pallens* assembly, made by the same NCBI pipeline as the reference annotation, offers a direct test of transfer loss for one form, and the Ensembl gene set of *Cx. perexiguus* shows what an independent annotation of a related genome recovers.

Results that do not depend on gene content hold up better. The phylogenomic analyses rest on single-copy orthologs present in all four forms, and synteny rests on the positions of shared genes. However, transfer errors affect both, and we have tried to measure this. The reference assembly performed best, with a protein-mode BUSCO score within one point of its genome-mode score.

### Genome Structure

Chromosome-scale synteny was conserved among the four forms. Most rearrangements were small. Those found in every comparison involving one assembly were most likely specific to that assembly, and their number followed assembly contiguity, from 31 in *molestus*, the assembly with the shortest contigs, to none in *pipiens*, the assembly with the longest. Rearrangements specific to one assembly could be real inversions, polymorphic in the sampled populations, or scaffolding errors. Their link to contiguity suggests many are scaffolding errors, but telling the two apart would require cytological or long-read evidence.

The 10 to 13% of shared genes placed on non-homologous chromosomes, concentrated near the middle of the chromosomes, most likely include placement errors from transfers in repetitive regions.

Repeats made up 57 to 59% of each assembly, consistent with the repeat content reported for the current *Cx. quinquefasciatus* assembly and about twice the estimate from the earlier, more fragmented assembly (Arensburger et al. 2010, Ryazansky et al. 2024), a gap that most likely reflects long-read assembly and de novo repeat libraries rather than biology.

### Limitations and Conclusions

Each form is represented by one assembly, so any difference between two forms includes variation within each. Where the genomes came from matters most for the gene flow results, because the extent of introgression between *Cx. pipiens* and *Cx. quinquefasciatus* differs by location (Cornel et al. 2003). The ABBA-BABA tests themselves cannot distinguish gene flow from ancestral population structure, and they do not show the direction of gene flow. The rooted tree and the main D statistics used the five taxon loci with intact ingroup models, about two-thirds of the total. These loci need not be a random genome sample, and residual transfer errors may remain. We transferred gene models without repairing broken coding sequences, which affected every gene-content analysis. CAFE ran on a tree with assumed branch lengths. The number of anvi’o gene clusters in each class depends on the minbit and MCL inflation settings; as run here, anvi’o separated broken models from their source genes more often than OrthoFinder did.

Within these limits, the four forms fall into two lineages, (*molestus*, *pipiens*) and (*pallens*, *quinquefasciatus*), that are closely related at the protein level. Their genomes carry an excess of derived alleles shared across the two lineages, between *molestus* and *quinquefasciatus* and between *pallens* and the *Cx. pipiens* lineage, consistent with gene flow within the complex and with the proposed hybrid origin of *pallens*. Apparent differences in gene content among the forms are mostly annotation transfer artifacts. Shared transfer errors can even mimic shared ancestry. Comparative genomics of this complex will need several genomes per form, each annotated independently, and the workflow and diagnostics described here offer a way to measure how much of a difference survives those checks.

## Author Contributions

Tyler Maire: Conceptualization, Data curation, Formal analysis, Investigation, Methodology, Project administration, Software, Visualization, Writing, original draft, Writing, review and editing. Erin Maley: Writing, review and editing. Theresa Miller: Writing, review and editing. Kyle Kosinski: Supervision, Writing, review and editing.

## Data Availability

The assemblies are available from NCBI under the accessions in Table 1, and the *Cx. perexiguus* gene set from Ensembl (Ruiz-López et al. 2026). The workflow, including the scripts that produce every table, figure, and reported value, is available at https://github.com/tylermaire/cx_pipiens_pangenome (release v3.0). The release and the larger outputs of the analysis (transferred annotations, protein and coding sequence sets, orthogroups, alignments, gene trees, CAFE outputs, anvi’o databases and validation alignments) are archived at Zenodo (Maire et al. 2026, https://doi.org/10.5281/zenodo.23460827). This article follows a preprint (bioRxiv, https://doi.org/10.64898/2026.07.25.740720).

## Supplementary Material

Supplementary data are available with this preprint as Supplementary Material. Supp. Tables S1 to S14 are provided in one workbook, with a title, caption, and list of source files for each table.

